# Sequential early-life viral infections modulate the microbiota and adaptive immune responses to systemic and mucosal vaccination

**DOI:** 10.1101/2023.08.31.555772

**Authors:** Yuhao Li, Jerome M. Molleston, Andrew H. Kim, Harshad Ingle, Somya Aggarwal, Lila S. Nolan, Ahmed O. Hassan, Lynne Foster, Michael S. Diamond, Megan T. Baldridge

## Abstract

Increasing evidence points to the microbial exposome as a critical factor in maturing and shaping the host immune system, thereby influencing responses to immune challenges such as infections or vaccines. To investigate the effect of early-life viral exposures on immune development and vaccine responses, we inoculated mice with six distinct viral pathogens in sequence beginning in the neonatal period, and then evaluated their immune signatures before and after intramuscular or intranasal vaccination against SARS-CoV-2. Sequential viral infection drove profound changes in all aspects of the immune system, including increasing circulating leukocytes, altering innate and adaptive immune cell lineages in tissues, and markedly influencing serum cytokine and total antibody levels. Beyond these immune responses changes, these exposures also modulated the composition of the endogenous intestinal microbiota. Although sequentially-infected mice exhibited increased systemic immune activation and T cell responses after intramuscular and intranasal SARS-CoV-2 immunization, we observed decreased vaccine-induced antibody responses in these animals. These results suggest that early-life viral exposures are sufficient to diminish antibody responses to vaccination in mice, and highlight their potential importance of considering prior microbial exposures when investigating vaccine responses.

## Introduction

House mice (*Mus musculus*) are regarded as a reliable and versatile model organism for biomedical research, because of their cost-effectiveness and inbred genetics, and availability of a wealth of reagents to characterize them phenotypically. However, over the past decade it has emerged that the limited natural microbial exposure in current specific pathogen-free (SPF) mouse facilities may have important influences on the relevance of research findings(1, 2). Despite having an immune system structured similarly to that of humans, with innate and adaptive components fulfilling similar roles, mice have often poorly replicated transcriptional responses and immune phenotypes seen in humans to the same stimuli, such that pre-clinical results fail to predict clinical trial outcomes(^3, 4^). Unlike humans and wild mice, which are both exposed to a wide variety of microbial influences, laboratory mice raised in SPF environments do not encounter the microbial stimuli found in nature, resulting in differential immune instruction and responses(^5–7^).

The recent paradigm shift in the understanding that immune phenotypes are potently regulated by these complex microbial exposures was brought about in part by various “dirty” mouse models. Cohousing of SPF mice with wild or pet shop-derived mice, rewilding by exposing SPF mice to a natural environment, and “wildling” models in which transferred lab mouse embryos are sired by wild mothers(6, 8–12), have provided critical comparisons between “exposed” and naïve SPF mice. These alternative “dirty” models reproduce natural microorganism exposures including bacteria, eukaryotic viruses, bacteriophages, fungi, helminths and/or mites(13, 14), and exhibit fundamental shifts in their immune systems, improving their ability to recapitulate essential aspects of human immunity(5, 6, 9). Co-housing laboratory mice results in the replication of immune phenotypes and transcriptional signatures found in wild mice or humans, including altering T cell differentiation and modulating susceptibility to new pathogens(6). Mice rewilded to an outdoors environment also exhibit notable maturation and differentiation within the T cell compartment, accompanied by heightened levels of circulating granulocytes, changes associated with an increase in gut colonization by fungal taxa(10). Wildling offspring mice also better phenocopy human immune responses to CD28-superagonist and anti-TNF-α treatments than SPF mice(9).

While these microbial transfer models have been useful to identify phenotypes governed by more varied microbial exposures, they are difficult to control due to the varied microbes present in any given pet shop or wild mouse or in the natural environment(2, 15). Furthermore, due to the potential danger for introducing hazardous microbes into animal facilities, experiments utilizing these models generally require specialized, dedicated facilities, which are not available to many investigators. An alternative approach is to instead employ sequential infection of SPF mice, in which animals are intentionally inoculated with a series of known pathogens. One such model, in which adult SPF mice were infected over time with three viruses and a helminth, demonstrated changes in gene expression that closely approximated pet shop mice as well as adult humans, and exhibited increased inflammatory cytokine responses but blunted antibody responses to yellow fever vaccine(^11^). This study suggested that transfer of the gut microbiota is not needed to “humanize” immune responses, but rather, it is the exposure to other pathogens that may be important. However, one challenge with the employment of this model was that the infection series was conducted in mice which had already reached adulthood, and was not completed until at least 15 weeks of age. Moreover, helminths themselves have their own microbiota(16, 17), which could independently impact immunity. Because the longitudinal microbiota was not profiled in these mice, the response of endogenous microbial communities to a sequential infection regimen remains unclear.

In an effort to develop a tractable sequential infection model for broad laboratory use, as well as to further describe the microbial and immune changes that result from sequential microbial exposures, we devised a virus-only sequential infection model beginning in early life that is completed by 6 weeks of age, allowing mice to be used for further experiments in a rapid and well-controlled manner. Using this model, we profiled changes in the immune system, identifying marked changes in immune cell populations as well as cytokine and antibody levels. We explored the antibody and T cell response to both intramuscular and intranasal SARS-CoV-2 vaccination in our sequentially-infected mice, and observed a blunted SARS-CoV-2-specific antibody response but enhanced effector T cell response. Finally, we profiled the intestinal microbiome and identified decreases in microbial diversity as well as changes in taxonomic composition associated with sequential viral infection. This tractable model enables the study of a more “humanized” immune system in mice in a rapid and consistent manner in standard animal facilities and implicates early-life virus infections as a regulator of the immune system of SPF mice.

## Results

### Sequential viral infection creates a durable pro-inflammatory host environment

We sought to develop an “immunologically-mature” murine model in a genetically defined background by utilizing a well-controlled series of microbial exposures, focused exclusively on viruses, that could be rapidly administered to permit further experimental intervention by the age of 10 weeks. Thus, we inoculated C57BL6/J mice with six distinct viruses - murine rotavirus strain EDIM-Cambridge (MRV, 10^4^ ID_50_ by oral gavage), murine gamma-herpesvirus 68 (MHV68, 10^5^ PFU intranasally), murine norovirus strain CR6 (MNV, 10^6^ PFU orally), influenza virus strain PR8 (IAV, 300 PFU intranasally), murine astrovirus (MAstV, infectious fecal filtrate by oral gavage), and coxsackievirus B3 (CVB3, 10^8^ PFU orally) - at 1-week intervals beginning when pups were 7 days old (**Figure 1A**). All viruses were administered at established inocula(11, 18–21) and viral shedding in the stool at expected intervals was confirmed for MNV and MAstV (**Fig S1A**). This sequential microbial exposure resulted in global immunological changes in the mice. Hematological analysis at 10 weeks of age revealed a pronounced leukocytosis on a background of preserved red blood cell and platelet counts (**Figure 1B, C**). Though the proportions of circulating lymphocytes, neutrophils, monocytes, eosinophils, and basophils were unchanged, their absolute counts were all significantly increased (**Figure 1C and Fig S1B)**. These phenotypes were not influenced by sex (**Fig S1C)**. Despite similar absolute red blood cell numbers, sequential infection was associated with decreased mean corpuscular hemoglobin (MCH) and mean corpuscular volume (MCV) and increased red cell distribution width (RDW-CV), changes potentially consistent with a chronic pro-inflammatory state (**Fig S1D**)(22).

**Figure 1.**
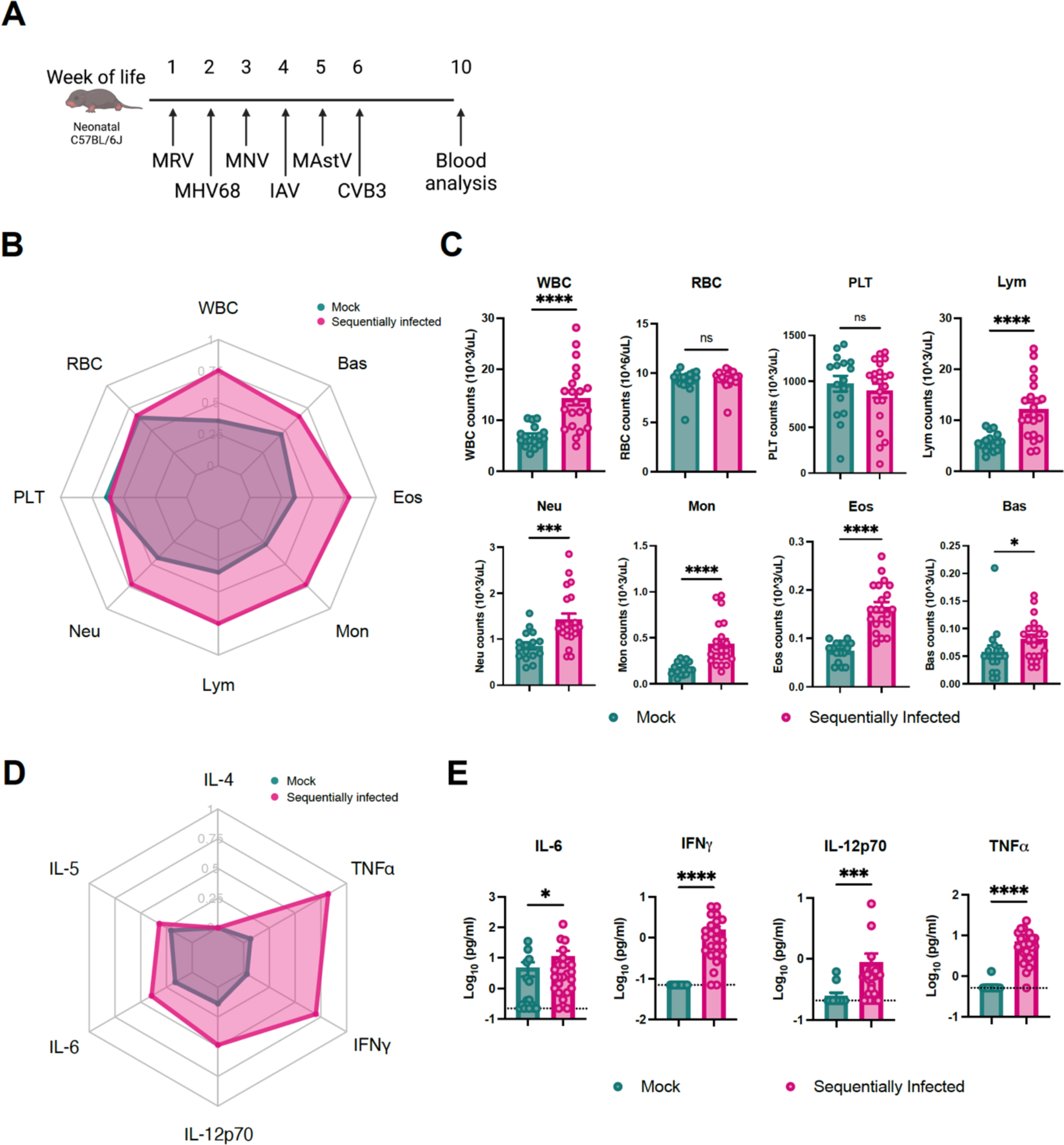
Sequential viral exposure creates an inflammatory host environment. (**A**) Abbreviated sequential infection regimen including murine rotavirus (MRV), murine gammaherpesvirus 68 (MHV68), murine norovirus CR6 (MNV), Influenza A Virus (IAV), murine astrovirus (MAstV), and coxsackievirus B3 (CVB3). Whole blood and indicated tissues of ten-week-old mock- and sequentially-infected C57BL/6J mice were collected for analysis. (**B, C**) Hematologic analysis of peripheral blood using the automatic blood cell counter Element HT5. Radar plot comparing the indicated blood cell populations in mock- (n = 16) and sequentially-infected mice (n = 21) (**B**). White blood cells (WBC), red blood cells (RBC), platelets (PLT), lymphocytes (Lym), neutrophils (Neu), monocytes (Mon), eosinophils (EOS), and basophils (BAS). (**D, E**) Serum was collected from mock- (n = 16) and sequentially-infected (n = 24) mice at the age of 10 weeks. The concentration of cytokines and chemokines was determined by Bioplex assay. Radar plot comparing the serum concentrations of indicated cytokines and chemokines in the indicated groups (**D**). Columns show median values and error bars represent the standard error of the mean, with dotted lines indicating the limit of detection (LOD) of the assays. Undetected cytokines were given a value of LOD. Statistical analysis was performed using unpaired Mann-Whitney test for (**C**) and (**E**): *p < 0.05; **p < 0.01; ***p < 0.001; ****p < 0.0001; ns, not significant.

In addition to circulating immune cells, we also profiled serum cytokine levels, which revealed substantial increases in pro-inflammatory cytokines interleukin (IL)-6 (mean fold change(FC) = 2.39, p= 0.043), IL-12p70 (FC = 3.53, p= 0.002), interferon (IFN)-γ (FC = 22.4, p= 0.001), and tumor necrosis factor (TNF)-α (FC = 12.4, p= 0.001) from sequentially-infected mice after 9 weeks of exposure (**Figure 1D, 1E and Fig S1E**). Together, these data demonstrate that sequential viral exposure increased multiple classes of circulating leukocytes and enhanced levels of inflammatory cytokines and chemokines in the serum.

### Sequential viral infection modulates the circulating and tissue-resident adaptive immune compartments of laboratory mice

A previous study showed that cohousing of SPF mice with pet shop mice does not alter the frequency and quantity of circulating T cells(5). In contrast, at 10 weeks of age, mice that had been sequentially-infected with viruses exhibited elevations in the total number of circulating CD4^+^ and CD8^+^ T cells (**Figure 2A and Fig S2A**), and this was associated with an enhanced proportion of CD8^+^ T cells (**Fig S3A**). To assess the impact of sequential viral infection on the differentiation state and distribution of CD8^+^ T cells, we monitored differentiation markers within the CD8^+^ T cell subset of peripheral blood mononuclear cells (PBMC). Peripheral blood antigen-experienced (CD44^hi^) CD8^+^ T cells were rapidly induced in sequentially-infected pups, with many displaying a differentiated effector memory (EM) phenotype (CD44^hi^CD62L^low^) (**Figure 2B and Fig S2A**). Consistent with the pattern observed in outbred/co-housed mice and humans,(5, 6, 23) sequentially-infected mice exhibited a higher proportion of activated cytotoxic (Grzb^+^) and terminal effector differentiated (CD62L^lo^, CXCR3^lo^ and KLRG1^+^) cells within the antigen-experienced (CD44^hi^) CD8^+^ T cell population (**Figure 2C and Fig S2B**). In contrast, the phenotypes of naïve (CD44^lo^) CD8^+^ T cells were indistinguishable between mock- and sequentially-infected mice. In addition, cell size (forward scatter, Fsc) and a T cell exhaustion marker (PD1) were equivalent between the two groups (**Figure 2C**).

**Figure 2.**
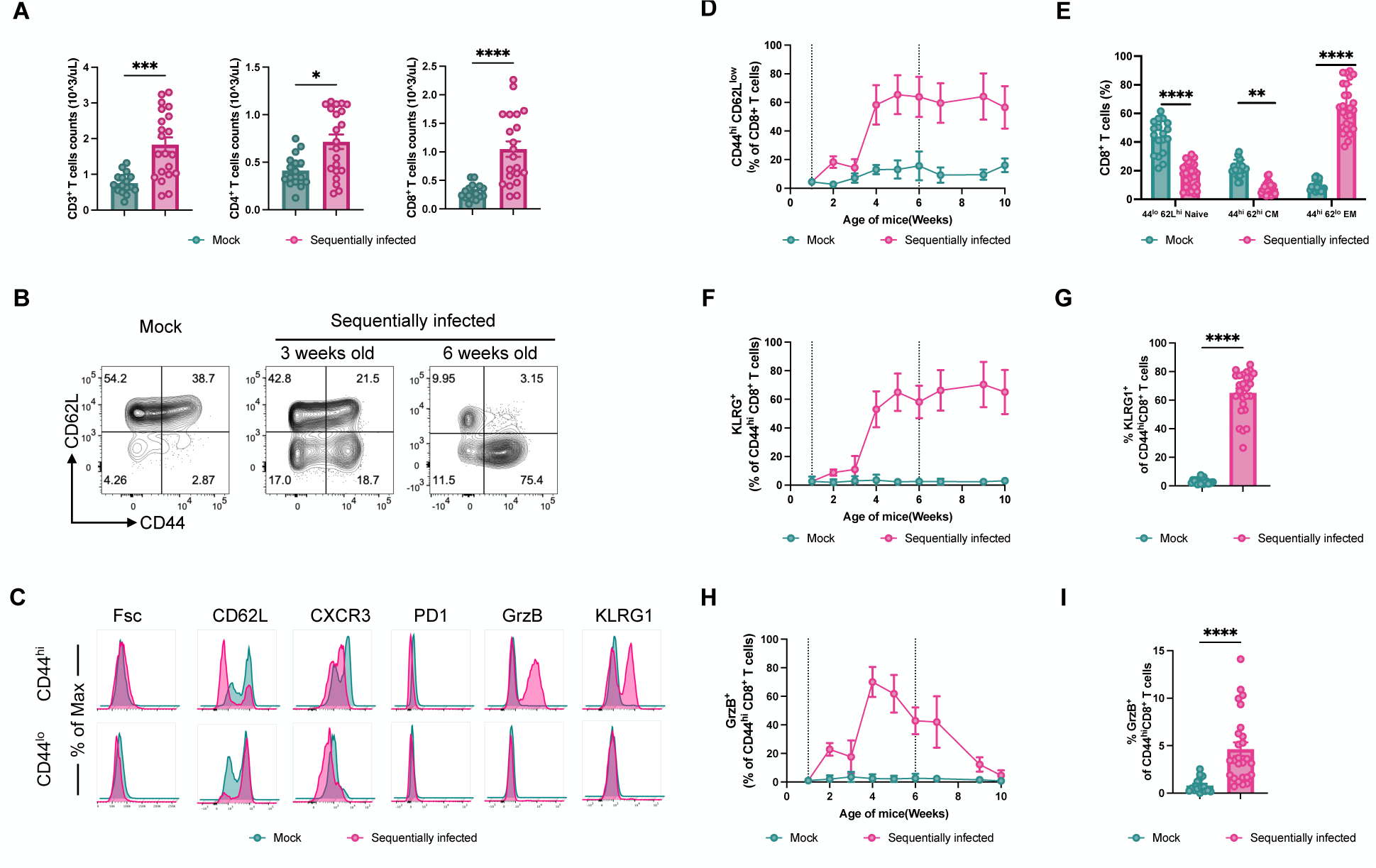
Sequentially infected mice have altered circulating immune cell compositions. (**A**) Peripheral blood T cell counts from mock- (n = 16) or sequentially-infected (n = 21) laboratory mice at the age of 10 weeks. (**B, C**) CD8^+^ T cell phenotypes in blood from mock- or sequentially-infected laboratory mice. Representative flow plots (n = 18 - 27) are shown (**B**). Phenotypes of CD44^lo^/CD62L^hi^ (naive) and CD44^hi^ (antigen-experienced) CD8^+^ T cells from PBMCs of mock- and sequentially-infected mice compared by flow cytometry (**C**). (**D, E**) Frequency of CD44^hi^ (antigen experienced) CD8^+^ T cells from PBMCs of mock-(n = 18) or sequentially-infected mice (n = 27) from four independent experiments (**D**). Comparison of CD44^lo^/CD62L^hi^ (Naive), CD44^hi^/CD62L^hi^ (antigen-experienced central memory, CM), and CD44^hi^/CD62L^lo^ (antigen-experienced effector memory, EM) CD8^+^ T cell between mock- and sequentially-infected mice at 10 weeks of age (**E**). (**F, G**) Frequency of KLRG^+^ CD44^hi^ CD8^+^ T cells from PBMCs of mock- (n = 18) or sequentially-infected mice (n = 27) from four independent experiments (**F**). Comparison of KLRG^+^ CD44^hi^ CD8^+^ T cell between mock- and sequentially infected mice at 10 weeks of age (**G**). (**H, I**) Frequency of GrzB^+^ CD44^hi^ CD8^+^ T cells from PBMCs of mock-mice (n = 18) or sequentially-infected mice (n = 27) from four independent experiments (**H**). Comparison of GrzB^+^ CD44^hi^ CD8^+^ T cell between mock- and sequentially-infected mice at 10 weeks of age (I). Columns show median values and error bars represent the standard error of the mean. Vertical dotted lines indicate the first and last viral inoculations. For (**A**), (**E**), (**G**), and (**I**) statistical analysis was performed using Mann-Whitney test: *p < 0.05; **p < 0.01; ***p < 0.001; ****p < 0.0001; ns, not significant.

Longitudinal analysis revealed that within six weeks of sequential infection, the proportion of CD44^hi^CD62L^low^ EM T cells within CD8^+^ PBMC increased from 5% to approximately 70%, and subsequently remained relatively stable for the duration of the study (**Figure 2B, 2D and Fig S3B**). At the age of 10 weeks, there were striking differences observed in the populations of naïve, central memory (CM), and terminally differentiated EM CD8^+^ T cells between the two groups of mice (**Figure 2E**). These changes in immune cell proportions were observed in mice of both sexes (**Fig S3C**). Moreover, the population of terminally differentiated EM cells characterized by KLRG1 expression also increased to over 60% in sequentially-infected mice. These KLRG1^+^ cells persisted in the peripheral blood throughout the study (**Figure 2F, 2G and Fig S3D**). Conversely, the population of GrzB+ effector differentiated memory CD8^+^ T cells, critical responders against viral infection, returned closer to baseline 4 weeks after the last viral exposure (**Figure 2H**). Nevertheless, sequentially-infected mice exhibited a higher abundance of GrzB^+^ cytotoxic T cells compared to mock-infected mice, indicating persistence of these effector cells despite the observed decrease from peak levels (**Figure 2I and Fig S3E**).

To gain a more comprehensive understanding of immune status beyond circulating phagocytes and T cells, we analyzed immune cell lineages in mucosal tissues of sequentially-infected mice (**Figure 3, Fig S4 and Fig S5**). In contrast to mock-infected mice, which displayed low numbers of CD8^+^ T cells in nonlymphoid tissues such as the lung and ileum, sequentially-infected mice had notable increases in the abundance of CD8^+^ T cells in these tissues (**Figure 3A, B**). Conversely, the number of mucosal regulatory T cells (Treg) and innate lymphoid cells (ILCs) were generally reduced after sequential infection (**Figure 3A, 3B and Fig S4A**). The mesenteric lymph node, the draining lymph node for the intestine, did not exhibit substantial changes in cell populations (**Fig S4B**).

**Figure 3.**
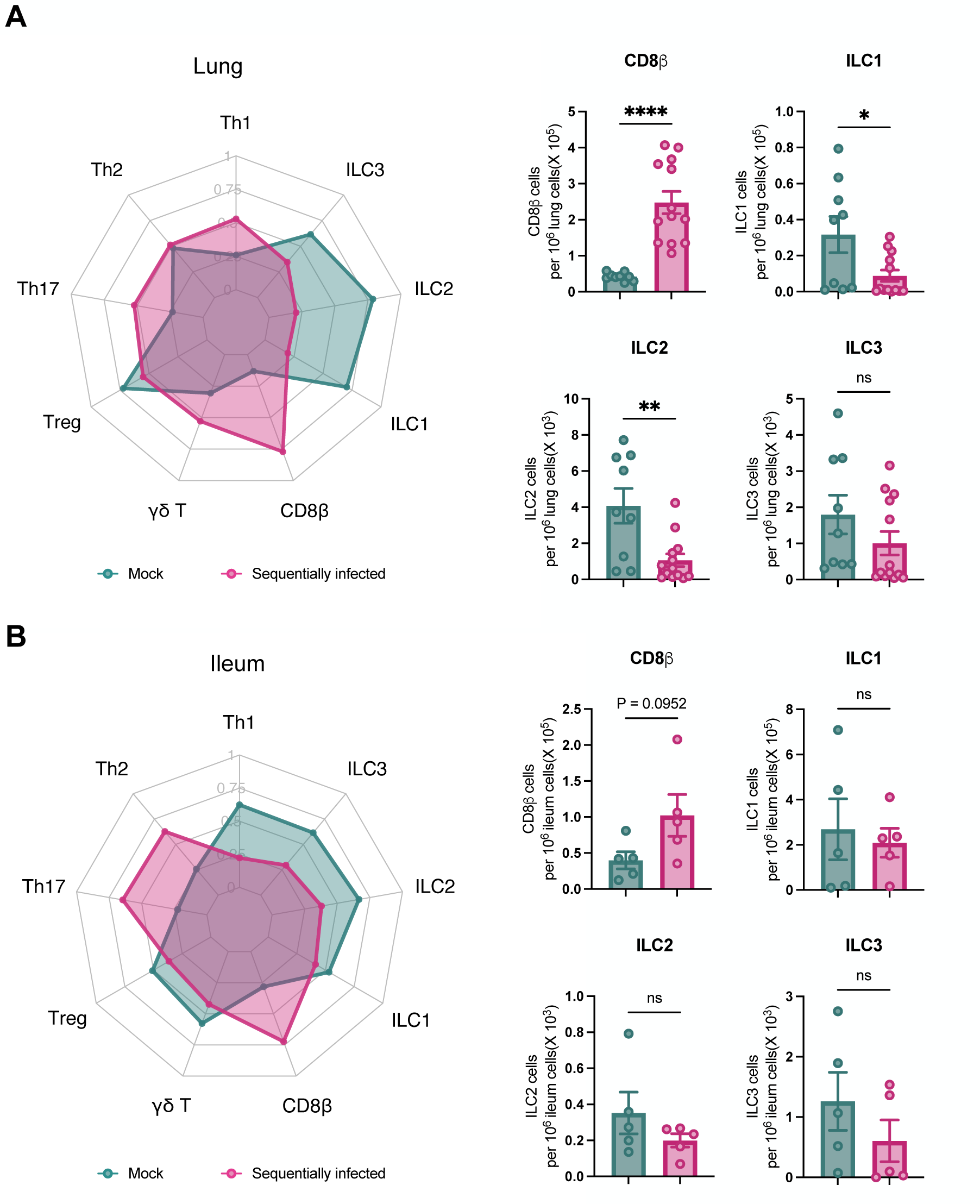
Sequential viral exposure shapes immune cell profiles in mucosal tissues. Cell types isolated and enumerated by flow cytometry from the lung (**A**) and ileum (**B**) of mock- and sequentially-infected 10 week-old mice (n=13/group); also shown in Fig. S4. ILC, innate lymphoid cells; T_h_, CD4 T helper cells; T_reg_, regulatory CD4 T cells; γδ T, gamma delta T cells. Radar plot comparing the indicated immune cell populations in mock- and sequentially-infected mice. Single-positive staining for T-bet, Gata3, and Rorγt were used to determine Th1, Th2, and Th17 lineages, respectively. Columns show median values and error bars represent the standard error of the mean. Statistical analysis was performed using unpaired Mann-Whitney test: *p < 0.05; **p < 0.01; ***p < 0.001; ****p < 0.0001; ns, not significant.

Beyond these changes in mucosal T cell populations, we next evaluated how sequential infection modulated antibody and B cell levels. We measured levels of total antibodies in serum, and found that all IgG subclasses as well as IgM and IgE, but not IgA, were significantly increased in sequentially-infected mice (**Figure 4A**). However, this increase in total antibodies in sequentially-infected mice was associated with significant reductions in splenic populations of immature B cells, B1 cells, and plasma cells, whereas germinal center (GC) B cells were increased (**Figure 4B and Fig S4C**). While the proportion of hematopoietic progenitors within the Lin^-^Sca1^+^c-Kit^+^ (LSK) fraction of the bone marrow (BM) was higher in sequentially-infected mice, frequencies of mature and immature B cells as well as B cell progenitors remained comparable (**Figure 4C and Fig S4C**). Taken together, these findings indicate that sequential infection augments both circulating and tissue-resident lymphocytes and antibody levels. Importantly, these phenotypes persisted for 4 weeks following the last exposure, indicating an enduring impact on immune homeostasis.

**Figure 4.**
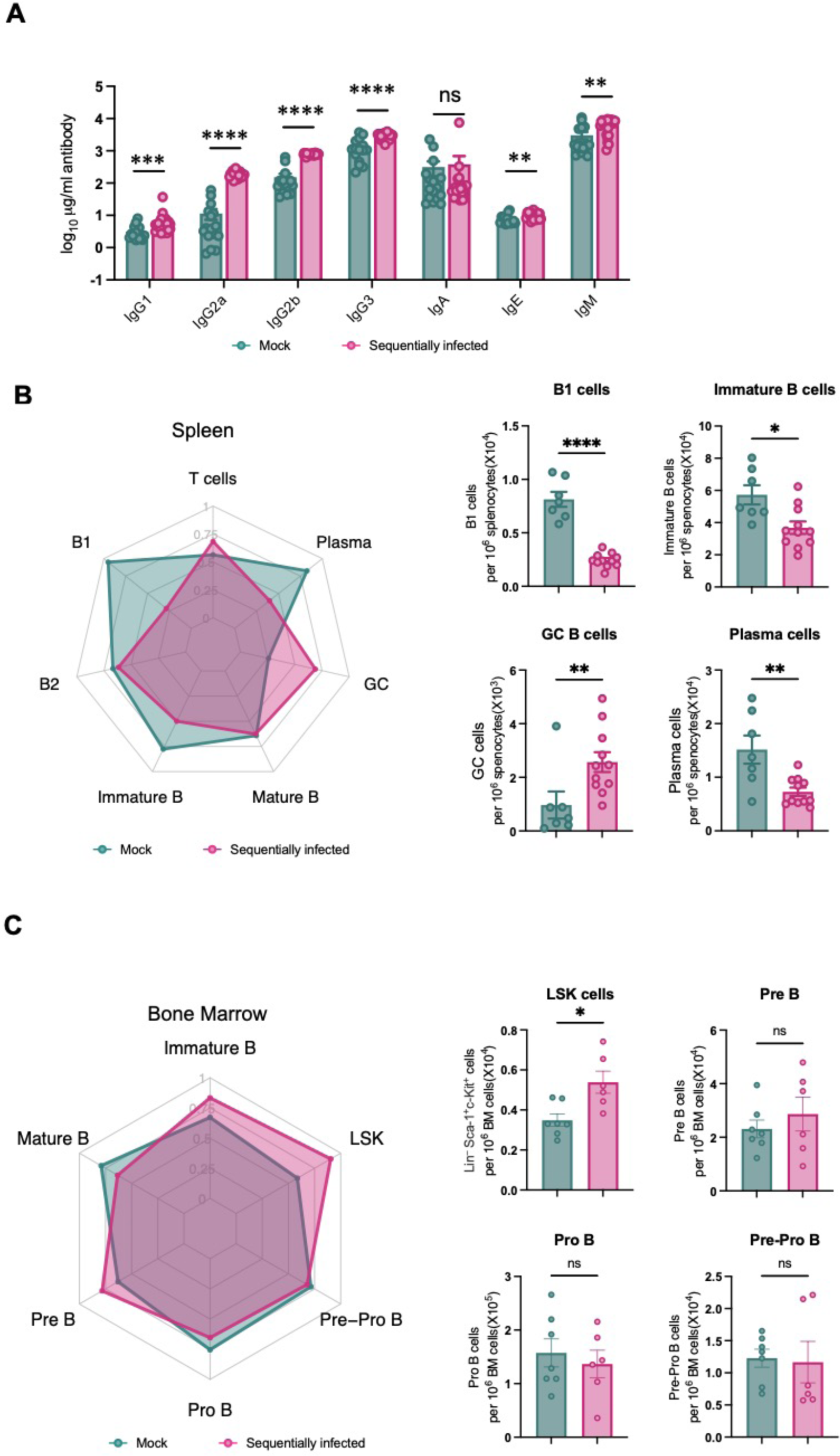
Sequential viral exposure modulates antibody levels and B cell lineages in tissues. (**A**) Total antibody concentrations in serum of mock- (n = 16) and sequentially-infected mice (n = 24) at 10 weeks of age. (**B, C**) Cell types isolated and enumerated by flow cytometry from the spleen (**B**) and bone marrow (BM) (**C**) of mock- and sequentially-infected mice. Radar plot comparing the indicated immune cell populations in mock- and sequentially-infected mice. Columns show median values and error bars represent the standard error of the mean. Statistical analysis was performed using unpaired Mann-Whitney test: *p < 0.05; **p < 0.01; ***p < 0.001; ****p < 0.0001; ns, not significant.

### Sequential viral infection modulates the intestinal microbiome

To evaluate the impact of sequential viral infection on the composition of major microbial niches, 16S ribosomal RNA (rRNA) gene profiling of the intestine and complete nasal turbinate (CNT) was surveilled from mock and sequentially infected mice at 9 weeks of age. A total of 43 fecal samples and 17 CNT samples were subjected to 16S rRNA amplicon sequencing, resulting in 324,493 and 23,088 reads, respectively. By employing dada2 for amplicon sequence variant (ASV) resolution and subsequent taxonomic assignment, we identified 250 distinct ASVs from the fecal samples and 107 distinct ASVs from the CNT samples (**Figure S6A and S6B**). Fecal samples obtained from sequentially-infected mice demonstrated a reduction in bacterial richness (Observed, the number of unique ASVs per sample) and diversity (Shannon, measure of both the number of unique ASVs and their evenness or distribution within sample) when compared to samples from mock-infected mice (**Figure 5A**, Observed: p = 0.0084; Shannon: p = 0.033). Intestinal bacterial evenness, assessed by Pielou’s evenness, was comparable between the two groups (p = 0.69). Conversely, no significant differences in alpha diversity parameters were observed in the samples collected from the CNT (**Figure 5A**; p = 0.39, 0.59, and 0.81 for Observed, Shannon, and Pielou indices, respectively). Principal coordinates analysis (PCoA) revealed distinct intestinal microbiota compositions between sequentially-infected mice and mock-infected mice (**Figure 5B**, PERMANOVA: p = 0.036). In contrast, no distinct clustering was observed in the CNT microbiota (**Figure 5B**, PERMANOVA: p = 0.68).

**Figure 5.**
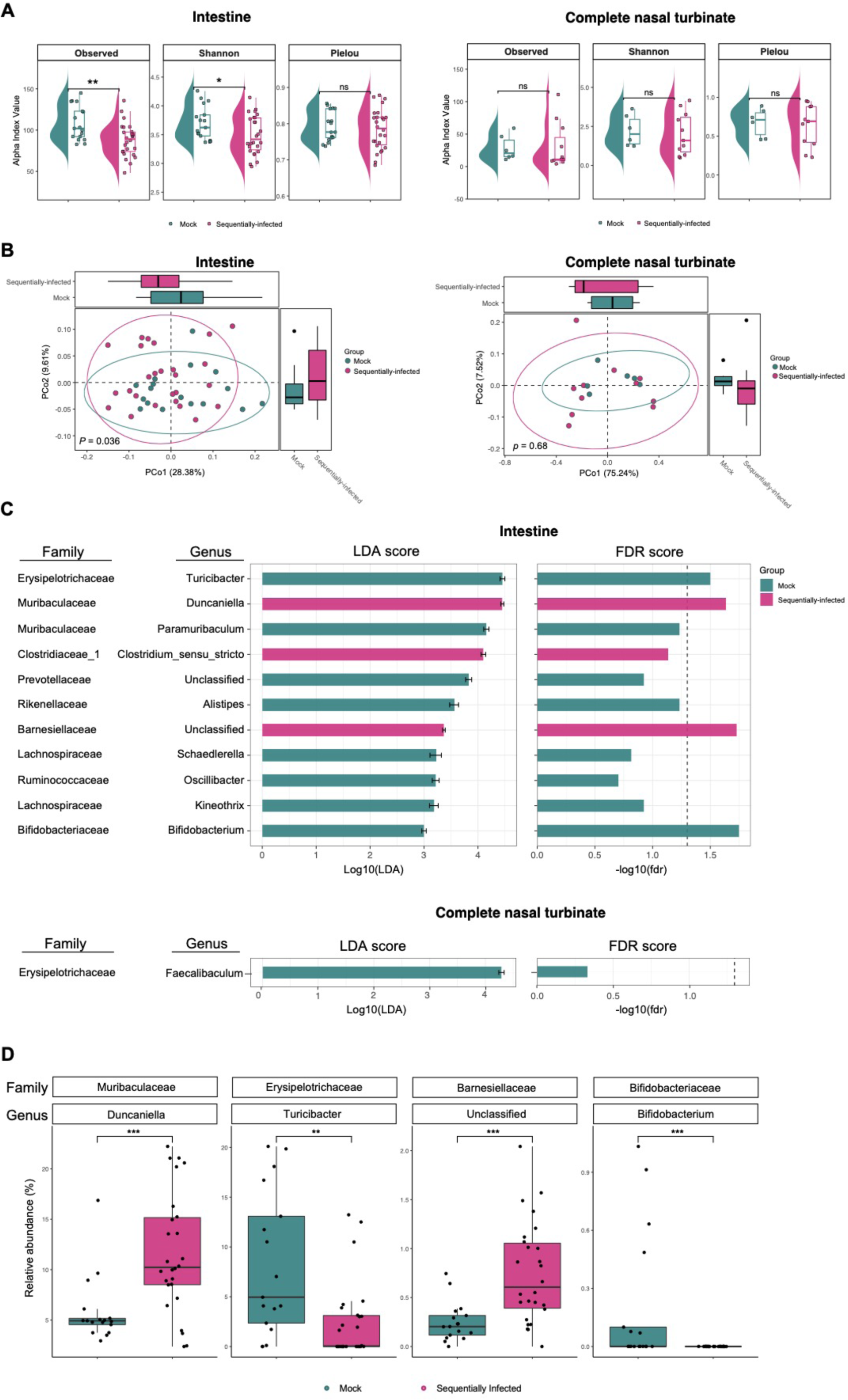
Sequential viral infection modulates the intestinal bacterial microbiome. The bacterial microbiome of mock- and sequentially-infected mice was profiled for fecal samples (mock-: n = 17, sequentially-infected: n = 26), and complete nasal turbinate tissues (mock-: n = 6, sequentially-infected: n = 11) using sequencing of the 16S rRNA gene. **(A)** Mean values and interquartile ranges of alpha diversity indices. Bacterial richness (Observed), Shannon diversity (Shannon), and Pielou’s evenness (Pielou) are shown. Mann-Whitney test: *p < 0.05; **p < 0.01; ***p < 0.001; ****p < 0.0001; ns, not significant. **(B)** Principal coordinates analysis (PCoA) using weighted UniFrac distances was performed. Differences between groups were assessed using PERMANOVA. **(C)** Linear discriminant analysis (LDA) effect size analysis was conducted to identify discriminant taxa between the intestinal and complete nasal turbinate microbiomes of mock- and sequentially-infected mice. In the left panel, the LDA effect size (with a 95% confidence interval) is displayed for taxa that were found to be distinct between groups. On the right panel, the false discovery rate (FDR) of each taxa is depicted, with vertical dotted line indicating an FDR of 0.05. **(D)** The relative abundance of discriminant taxa with low FDR identified from the intestinal microbiota at the genus level. Analyzed by Wilcoxon-Mann-Whitney test: **p < 0.01; ***p < 0.001.

To further explore potential differences in the bacterial communities between mock- and sequentially-infected groups, we performed Kruskal-Wallis rank sum tests and linear discriminant analysis (LDA) based on ASV abundances. These analyses revealed differentially-abundant taxa between the two groups (**Figure 5C**). Specifically, the relative abundances of *Duncaniella* (p = 0.001, fdr = 0.023), *Clostridium sensu stricto* (p = 0.011, fdr = 0.007), and an unclassified genus belonging to the *Barnesiellaceae* family (p = 0.0007, fdr = 0.019) were higher in the intestinal microbiota of sequentially-infected mice (**Figure 5C and S6C**). Conversely, the intestinal microbiota of sequentially-infected mice exhibited reduced relative abundances of *Paramuribaculum* (p = 0.007, fdr = 0.058), *Turicibacter* (p = 0.002, fdr = 0.031), *Alistipes* (p = 0.007, fdr = 0.058), *Oscillibacter* (p = 0.048, fdr = 0.197), *Schaedlerella* (p = 0.035, fdr = 0.153), *Kineothrix* (p = 0.024, fdr = 0.119), *Bifidobacterium* (p = 0.0004, fdr = 0.018), and an unclassified genus belonging to the *Prevotellaceae* family (p = 0.023, fdr = 0.119) (**Figure 5D and S6C**). In the CNT, only the relative abundance of *Faecalibaculum* (p = 0.012, fdr = 0.47) was decreased in sequentially-infected mice (**Figure 5D and S6D**). These findings indicate that our sequential viral exposure regimen modulates the composition of the intestinal microbiota, with minimal effects on the CNT microbiota, of SPF mice.

### Sequential viral infection limits antibody responses and enhances T cell responses to a SARS-CoV-2 vaccine administered by intramuscular or intranasal routes

We next sought to functionally evaluate how early-life microbial exposures influenced responses to a subsequent immune challenge. We used a recently-developed chimpanzee adenovirus-vectored vaccine encoding a prefusion stabilized spike protein (ChAd-SARS-CoV-2-S)(24) that is currently deployed in India (iNCOVACC®) and can be readily administered by intramuscular (IM) or intranasal (IN) routes, thereby permitting characterization of effects on systemic and mucosal immunity. At 10 weeks of age, we administered a single 10^10^ viral particle dose of ChAd-SARS-CoV-2-S to mice that had been sequentially-infected or mock controls. Serum samples were collected 4 weeks after immunization, and IgG and IgA responses against purified S proteins were evaluated by enzyme-linked immunosorbent assay (ELISA). While the ChAd-SARS-CoV-2-S vaccine elicited robust levels of S-specific IgG in both mock- and sequentially-infected mice, the S-specific IgG responses in sequentially-infected mice were lower (IM: FC=0.53, p=0.02; IN: FC=0.55, p=0.005) regardless of route of vaccination (**Figure 6A and 6B**). Consistent with a previous report(24), we observed that IM administration of the ChAd-SARS-CoV-2-S vaccine did not induce S-specific serum IgA in either group (**Figure 6A**). In contrast, IN immunization with ChAd-SARS-CoV-2-S elicited S-specific IgA in both groups, but this too was reduced in sequentially-infected mice (**Figure 6B**).

**Figure 6.**
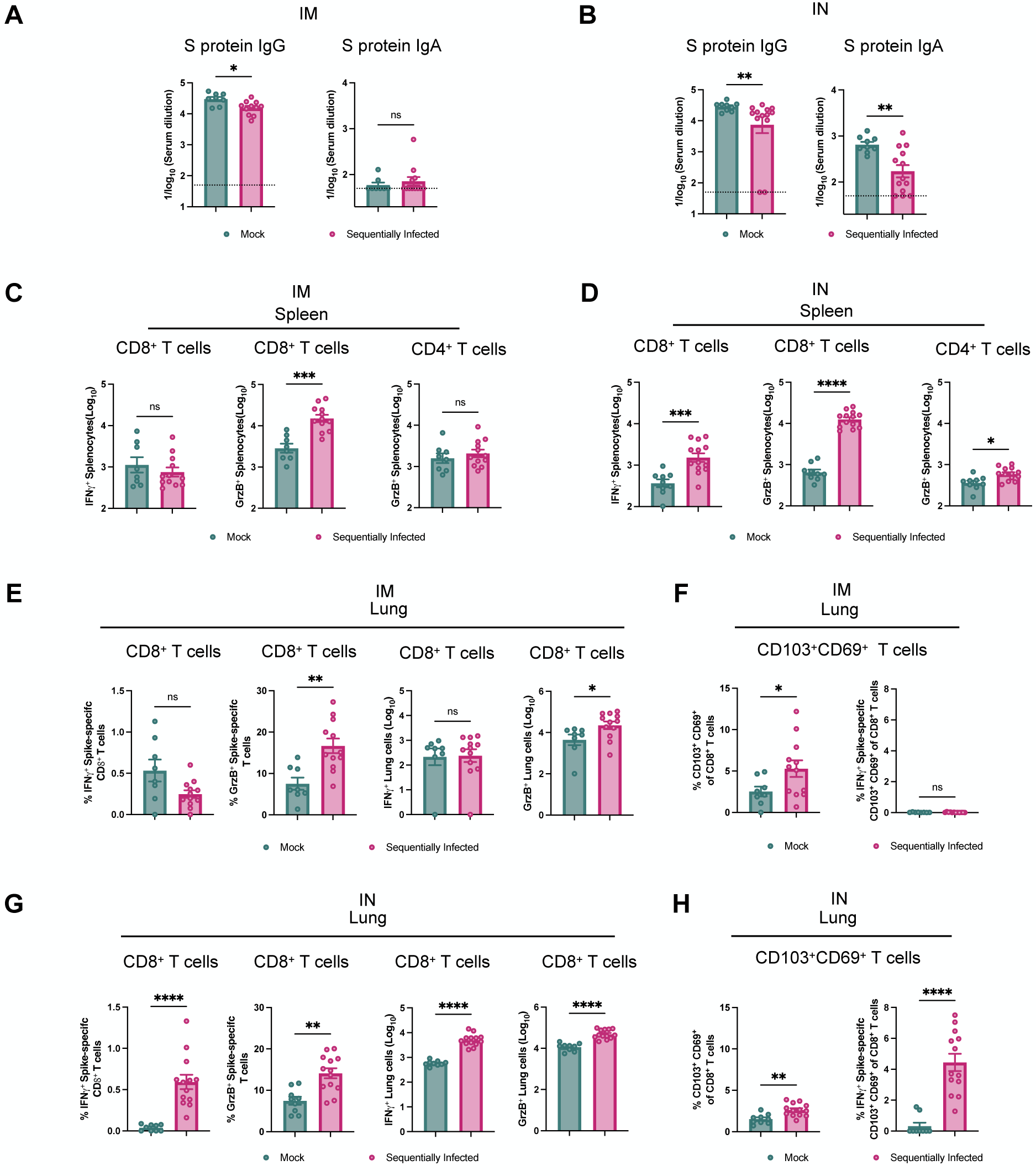
Sequential viral infection is associated with reduced antibody response to intramuscular and intranasal ChAd-SARS-CoV-2 vaccination. (**A, B**) Antibody responses in sera of immunized mice at day 28 after priming were evaluated. Anti-Spike IgG and IgA levels were measured by ELISA in intramuscular (**A**) or intranasal (**C**) vaccination studies. Data are pooled from two experiments (n = 8–13). (**C, D**) Cell-mediated immune responses were analyzed at day 35 post-immunization after re-stimulation with an S protein-peptide pool. Splenocytes were assayed for IFN-γ and GrzB expression in CD8^+^ T cells and GrzB only in CD4^+^ T cells by flow cytometry. Cell counts of positive splenocytes in intramuscular (**C**) or intranasal (**D**) vaccination studies (n = 8-13). (**G - H**) Lung CD8^+^ T cells from intramuscularly (**E, F**) or intranasally (**G, H**) vaccinated mice were assayed for IFN-γ and GrzB expression by flow cytometry after re-stimulation with an S protein peptide pool. Lung CD8^+^ T cells were phenotyped for expression of CD103 and CD69 after intramuscular (**F**) or intranasal (**H**) vaccination. Columns show median values and error bars represent the standard error of the mean. Dotted lines indicate the LOD of the assays. Mann-Whitney test: *p < 0.05; **p < 0.01; ***p < 0.001; ****p < 0.0001; ns, not significant.

To evaluate vaccine-induced T cell responses, splenocytes and lung cells were harvested 5 weeks post-vaccination and stimulated *ex vivo* with a pool of 253 overlapping 15-mer S peptides pool as previously described(25). Following *ex vivo* peptide re-stimulation, we observed a significantly higher number of splenic CD8^+^ T cells expressing GrzB in sequentially-infected mice that received IM immunization (**Figure 6C and S7**). This finding suggests that microbial exposure enhances the cytotoxic potential of CD8^+^ T cells, potentially enabling them to better eliminate infected cells. However, there were no significant increases observed in the numbers of CD8^+^ T cells expressing IFN-γ or CD4^+^ T cells expressing GrzB, indicating that sequential infection may not globally affect the polyfunctionality of these cellular subsets after IM vaccination. In contrast, after IN immunization, we observed an increase in the frequency of splenic IFN-γ-expressing CD8^+^ T cells, as well as GrzB-expressing CD4^+^ and CD8^+^ T cells, in sequentially-infected mice (**Figure 6D**). This suggests that sequential microbial exposure enhances the activation and effector functions of the T cell subsets, particularly following IN vaccination.

In evaluating cells from the lung, we observed a significant increase in the frequency of GrzB-producing CD8^+^ T cells in sequentially-infected mice that received IM vaccination (**Figure 6E**). However, the frequency of IFN-γ- secreting, antigen-specific CD103^+^CD69^+^CD8^+^ T cells within this subset remained relatively low (**Figure 6F**). This indicates that although the overall presence of lung-resident memory T cells was augmented, their capacity to produce IFN-γ in response to antigen stimulation was not altered for IM vaccination. However, upon *ex vivo* re-stimulation of lung cells from sequentially-infected mice following IN immunization, a notable and significant increase in the population of T cells producing both IFN-γ and GrzB was observed (**Figure 6G**). Thus, although the baseline frequency of IFN-γ-secreting, antigen-specific resident memory T cells was initially low, their functionality and capacity to produce cytokines were enhanced by sequential infection. Furthermore, the increase in the frequency of IFN-γ-secreting, antigen-specific CD103^+^CD69^+^CD8^+^ T cells in the lung (**Figure 6H**), supports the notion that sequential infection promotes the generation of tissue-resident memory T cells with enhanced cytokine secretion capacity. Overall, these findings indicate that repeated viral exposures impact responses to vaccination by blunting vaccine-specific antibody responses and potentially enhancing antigen-specific T cell responses particularly following mucosal exposure. These findings highlight the importance of considering exposure history in evaluating pre-clinical interventions, as well as the immunological consequences of early-life viral infections.

## Discussion

While the efficacy of vaccines relies on antibody- and immune cell-mediated responses, the magnitude of these responses varies across individuals(26–28). Those residing in high-income countries (HICs) often display a more potent immune response to vaccination than those in low-income and middle-income countries (LMICs)(29). These distinctions in immunological reactions have been noted for vaccines against influenza, yellow fever, and Ebola(30–32). Differential microbiota composition and diversity between HICs and LMICs, as well as differential microbial exposure histories, have been implicated as potentially driving these differential vaccine responses(29, 33). Perturbation of the gut microbiome with antibiotics as well as viral co-infections can have important effects on human vaccine responses(34–37). The initial state of the intestinal microbiota has also been linked to the immunogenicity of and adverse effects associated with SARS-CoV-2 vaccines(38). Furthermore, the utilization of probiotics such as *Lactobacillus* and *Bifidobacterium*, has exhibited potential in augmenting immune responses to vaccines(39–41). However, the precise identity of intestinal microbes or prior infections that may influence vaccine effectiveness, as well as the mechanisms underlying this influence, remain enigmatic.

Rodent models with heightened microbial exposures, either via cohousing with “dirty” mice or via sequential infection,(13) have been shown to exhibit diminished humoral responses to vaccination(8, 11). In this study, we investigated the effects of sequential viral infections in early life on the immune system, vaccine responses, and microbiota of laboratory mice. Six distinct viral exposures resulted in profound changes to the immune system of the mice, including changes in the number and differentiation of white blood cells, increased pro-inflammatory cytokines and total antibodies in serum, and altered peripheral blood antigen-experienced CD8^+^ T cells. The differentiation of CD8^+^ T cells was rapidly induced and antigen-experienced CD8^+^ T cells increased to approximately 70% of peripheral blood mononuclear cells. Single-dose immunization of SARS-CoV-2 stabilized S-protein based vaccine of sequentially-infected mice resulted in enhanced antigen-specific CD8^+^ T cell responses but decreased antibody responses for both IM and IN routes. In addition, we observed changes to the intestinal microbiota associated with sequential viral infection, despite the absence of any direct modulation of intestinal bacterial populations.

Our findings parallel those observed in immune characterization of other “dirty” mouse models(6). As in our model, SPF mice cohoused with pet shop mice display decreased antibody responses to vaccination^7^. The elevated cytokines seen in our sequentially infected mice are similar to those seen in “wildling” models as well as to a prior sequential infection model utilizing viruses and a helminth^8,9^. Our study suggests that many major immune phenotypes observed in “dirty” mouse models could be driven by transmission of viral infections including astroviruses and enteroviruses, which has been recently documented as an important consequence of cohousing with pet shop mice(15). Indeed, our observation of an altered intestinal bacterial microbiota secondary to sequential infection also supports that either viral infection itself or the indirect effects of induced immune responses may contribute to modulation of endogenous microbial communities observed in other “dirty” models.

Compared to these other models, sequential viral infection in early life has the advantage of tractability, with a reduced set of controllable exposures that can be performed in a standard SPF mouse facility. Compared to the longer timecourse of prior sequential infection models(11), this model is expeditious, with the infection series complete by 6 weeks of life. This model thus permits specific dissection of the microbial factors and host pathways that contribute to these immune phenotypes. While the variable exposures achieved via cohousing of SPF mice with wild or pet shop mice may better model the breadth of immune exposures experienced by humans(14), intentional exposures that appropriately mimic the “average” exposures experienced by humans in HICs or LMICs also could inform our understanding of the diversity of human immune responses.

There remain several important limitations to this study and unanswered questions that we aim to address in the future. While the current model uses six viruses to mimic a wide range of viral exposures, it is still unknown which of the viruses are essential to immune maturation, or how critical the relative sequence of infections may be. Two of the viruses used in this study, MHV68 and MNV, establish chronic infection in mice, while the others are self-limited; it is possible that the immune exposure of acute infection is the key event, or that ongoing immune stimulation associated with chronic and latent viral infection may maintain the immune activation seen in this study(42, 43). It is possible that a further reduction in microbial stimuli may be sufficient to mature the immune system. Furthermore, the importance of the bacterial microbial changes in contributing to the immune maturation we observed is not yet clear. Further investigation is needed to determine whether these changes are a major factor in the altered immune system composition and responses of sequentially-infected mice.

Overall, the results of this study indicate that sequential viral infections modulate the microbiota and lead to changes in the immune system that dampen specific adaptive responses to systemic and mucosal vaccination. This study highlights the importance of early-life microbial exposure and its impact on the immune system and gut microbiota. Sequential infection provides a powerful model for a matured immune system that can be readily leveraged for immunology, virology, and vaccine studies.

## Materials and Methods

### Mice and sequential infections

WT C57BL/6J (stock no. 000664) mice were purchased from Jackson Laboratories and maintained at Washington University School of Medicine under specific-pathogen-free conditions according to University guidelines. Timed matings were performed, with males subsequently removed from breeding cages. Mice were initially infected at 7 days of life, and subsequent infections and analyses were performed as indicated, with mice weaned from dams at 21 days of life. Weaned mice were then co-housed with up to five mice of the same sex per cage with autoclaved standard chow pellets and water provided *ad libitum*.

Neonatal mice were inoculated with six distinct viral pathogens - murine rotavirus strain EDIM-Cambridge (MRV, 10^4^ ID_50_/mouse), murine gamma-herpesvirus 68 (MHV68, 10^5^ PFU/mouse), murine norovirus strain CR6 (MNV, 10^6^ PFU/mouse), influenza virus strain PR8 (IAV, 300 PFU/mouse), murine astrovirus (MAstV), and coxsackievirus B3 (CVB3, 10^8^ PFU/mouse) - at 1-week intervals beginning when pups were at 1 week of life. Blood and fecal samples were collected for virological, immunological, and microbiota analyses. Four weeks after the last viral exposure, 10-week-old animals were immunized with a single dose of 10^10^ viral particles (vp) of ChAd-SARS-CoV-2-S in 50 µl PBS via intramuscular injection in the hind leg or via intranasal inoculation(24, 44). Animals were euthanized at 5 weeks post-vaccination, and tissues were harvested for virological, immunological, and pathological analyses. Animal studies were carried out in accordance with the recommendations in the Guide for the Care and Use of Laboratory Animals of the National Institutes of Health. The protocols were approved by the Institutional Animal Care and Use Committee at the Washington University School of Medicine (22–0140).

### Blood analysis for hematology

To evaluate the complete blood counts (CBC) from the mock- and sequentially-infected mice, blood was collected via the submandibular vein into EDTA-coated tubes and promptly analyzed using the Heska Element HT5 hematology analyzer (Barrie, Ontario, Canada).

### ELISA

Purified SARS-CoV-2 S protein was coated onto 96-well Maxisorp clear plates at 2 mg/mL in 50 mM Na_2_CO_3_ pH 9.6 (70 µL) overnight at 4°C. Coating buffers were aspirated, and wells were blocked with 200 µL 1% BSA PBST (Blocking buffer) for 1 h(hour) at 37°C. The blocked plates then were ready for use after three rounds of PBST washing. Heat-inactivated serum samples were diluted in blocking buffer, and 50 µL of respective serum dilutions were used for the assay. Diluted serum samples were incubated in the blocked ELISA plates for 1 h at room temperature. The ELISA plates were again washed thrice in PBST, followed by the addition of 50 µL of anti-mouse IgG-HRP (, 1:2000 in PBST) or anti-mouse IgG-HRP (, 1:2000 in PBST). Plates were incubated at room temperature for 1 h, washed thrice in PBST, and then 1:5000 dilution of streptavidin-HRP (Thermo-Fisher) was added to wells. Following a 1 h incubation at room temperature, plates were washed thrice with PBST, and 50 mL of 1-Step Ultra TMB-ELISA was added (ThermoFisher Cat. #34028). Following a 12 to 15-min incubation, reactions were stopped with 50 mL of 2 M sulfuric acid. The absorbance of each well at 450 nm was read (Synergy H1) within 2 min of the addition of sulfuric acid. Optical density (450 nm) measurements were determined using a microplate reader (Bio-Rad). The endpoint serum dilution was calculated with curve fit analysis of optical density (OD) values for serially diluted sera with a cut-off value set to three times the background signal.

### Flow cytometry characterization

Peripheral blood leukocytes, splenocytes, or tissue cells were evaluated by flow cytometry. RBCs were removed from heparinized whole blood or single-cell suspensions using RBC lysis buffer (BioLegend). Isolated mouse cells were surface-stained with antibodies against CD3 (BD Biosciences, 145-2C11,1:200), CD45(Biolegend, 30F-11,1:200), CD11b(Biolegend, M1/70,1:200), CD11c(Biolegend, N418,1:200), GR1(Biolegend, RB6-8C5,1:200), Ly6G(Biolegend, 1A8,1:200), Ly6C(Biolegend, AL21, 1:200), MHC II (Ia-Ie)(Biolegend, M5/114.15.2, 1:200), CD8α(Biolegend, 53-6.7,1:400), CD4(Biolegend, RM4-5, 1:400), CD62L (Biolegend, MEL-14, 1:200), CD44(Biolegend, IM7, 1:200), CD69(Biolegend, H1.2F3, 1:200), CD103(Biolegend, M290, 1:200), PD-1(Biolegend, RMP1-30, 1:200), KLRG1(Biolegend, 2F1, 1:200), CXCR3(Biolegend, CXCR3-173, 1:200), CD127(Biolegend, SB/199, 1:200), TCR γ/δ (Biolegend, GL3, 1:200), F4/80(BD Biosciences, Cl-A3-1, 1:200), GrzB(Invitrogen, GRB04, 1:200), CD43(Biolegend, 1B11,1:200), IgM(Biolegend, RMM-1,1:200), IgD (Biolegend, 11-26c.2a,1:200,), Ter119(Biolegend, TER-119,1:200), Gr-1(Biolegend, RB6-8C5,1:400), CD19(Biolegend, 6D5,1:200,), B220(Biolegend, RA3-6B2,1:200), Sca-1(Biolegend, D7,1:200,), c-Kit(eBioscience, 2B8,1:100,). Cell viability was determined using LIVE/DEAD™ Fixable Aqua Dead Cell Stain Kit (Invitrogen). Intracellular staining for transcription factors was performed using the eBioscience™ Foxp3 / Transcription Factor Staining Buffer Set (Invitrogen) with antibodies against granzyme B (QA16A02), Foxp3 (FJK-16s), T-bet (4B10), Eomes (Dan11mag), Gata3 (L50-823), Rorγt (Q31-378) following manufacturer’s guidelines. Single positive staining for T-bet, Gata3, and Rorγt were used to determine Th1, Th2, and Th17 lineages, respectively. The stained samples were acquired using BD X-20 cytometer and analyzed with FlowJo X 10.0 software.

### Peptide restimulation and intracellular cytokine staining

Splenocytes from intramuscularly vaccinated mice were incubated in culture with a pool of 253 overlapping 15-mer SARS-CoV-2 S peptides(25) for 12 h at 37°C before a 4 h treatment with brefeldin A (BioLegend, 420601). Following blocking with an anti-FcγR antibody (BioLegend, clone 93), cells were stained on ice with CD45 (BD BioSciences, 30-F11, 1:400); CD44, CD4, CD8b, and CD19 (BioLegend, 1:200, respectively), and Fixable Aqua Dead Cell Stain (Invitrogen, L34966). Stained cells were fixed and permeabilized with the Foxp3/Transcription Factor Staining Buffer Set (eBiosciences, 00-5523). Subsequently, intracellular staining was performed with anti-IFN-γ (BD Biosciences, XMG1.2, 1:200), anti-TNF-α (BioLegend, MP6-XT22, 1:200), and anti-GrB (Invitrogen, GRB04, 1:200). Lungs from immunized mice were harvested and digested for 1 h at 37°C in digestion buffer consisting of RPMI media supplemented with Collagenase (Sigma, 2 mg/ml) and DNase I (Sigma, 0.05 mg/ml). Lung cells were incubated at 37C with the pool of 253 overlapping 15-mer SARS-CoV-2 S peptides described above in the presence of brefeldin A for 5 h at 37C. Lung cells then were stained as described above except no CD19 staining was included, and CD103-FITC and CD69-BV711 (BioLegend clones, 2E7, and, H1.2F3, respectively) were added. Analysis was performed on a BD X-20 cytometer, using FlowJo X 10.0 software.

### Measurement of serum cytokines, chemokines, and antibodies

Serum cytokines, chemokines, and antibodies were quantitated with the multiplex immunoassay Th1/Th2 panel 6plx (IL-4, IL-5, IL-6, IFN-γ, IL-12p70, and TNF-α) or mouse Antibody Isotyping Panel 7plx (IgG1, IgG2a, IgG2b, IgG3, IgA, IgE, and IgM) using a Luminex 200 with Bio-plex Manager Software 5.0.

### 16S rRNA gene Illumina sequencing and analysis

Phenol:chloroform-extracted DNA from fecal pellets was used for both 16S rRNA gene qPCR and sequencing. SYBR green qPCR for the 16S rRNA gene was performed with 515F (5’-GTGCCAGCMGCCGCGGTAA-3’) and 805R (5’-GACTACCAGGGTATCTAATCC-3’) primers to detect the V4 hypervariable region.

Primer selection and PCRs were performed as described previously (45). Briefly, each sample was amplified in triplicate with Golay-barcoded primers specific for the V4 region (F515/R806), combined, and confirmed by gel electrophoresis. PCR reactions contained 18.8μL RNase/DNase-free water, 2.5μL 10X High Fidelity PCR Buffer (Invitrogen), 0.5μL 10 mM dNTPs, 1μL 50 mM MgSO4, 0.5μL each of the forward and reverse primers (10 μM final concentration), 0.1μL Platinum High Fidelity Taq (Invitrogen) and 1.0μL genomic DNA. Reactions were held at 94°C for 2 min to denature the DNA, with amplification proceeding for 26 cycles at 94°C for 15s, 50°C for 30s, and 68°C for 30s; a final extension of 2 min at 68°C was added to ensure complete amplification. Amplicons were pooled and purified with 0.6x Agencourt Ampure XP beads (Beckman-Coulter) according to the manufacturer’s instructions. The final pooled samples, along with aliquots of the three sequencing primers, were sent to the Center for Genome Sciences (Washington University School of Medicine) for sequencing by the 2×250bp protocol with the Illumina MiSeq platform.

Read quality control and the resolution of amplicon sequence variants (ASVs) were performed with the dada2 R package (46). ASVs that were not assigned to the kingdom Bacteria were filtered out. The remaining reads were assigned taxonomy using the Ribosomal Database Project (RDP trainset 16/release 11.5) 16S rRNA gene sequence database (47). Ecological analyses, such as alpha-diversity (richness, Faith’s phylogenetic diversity) and beta-diversity analyses (unweighted and weighted UniFrac distances), were performed using PhyloSeq and additional R packages(48), and differentially abundant taxa between sample groups were identified by performing pairwise comparisons using LEfSe and MicrobiotaProcess package(49), (50). 16S rRNA gene sequencing data have been uploaded to the European Nucleotide Archive (accession no. PRJEB65723).

### Statistical analysis

Statistical significance was assigned when P values were <0.05 using Prism Version 8 (GraphPad). Tests, number of animals (n), median values, and statistical comparison groups are indicated in the figure legends. Radar charts were plotted using an R library(51).

## Data and materials availability

The data from this study are tabulated in the main paper and supplementary materials. All reagents are available from M.T.B. under a material transfer agreement with Washington University.

## Acknowledgments

We acknowledge all members of the Baldridge laboratory for helpful discussions.

## Author contributions

Y.L., J.M.M., A.H.K., H.I., S.A., L.S.N. and L.F. performed the experiments. Y.L. and M.T.B. analyzed the results. A.O.H. and M.S.D. provided the ChAd-SARS-CoV-2 vaccine and helped with experimental design. Y.L., J.M.M., and M.T.B. designed the project. Y.L., J.M.M. and M.T.B. wrote the manuscript. All authors read and edited the manuscript.

