## Supplementary Material for "Sequential early-life viral infections modulate the microbiota and adaptive immune responses to systemic and mucosal vaccination"

Figures S1-S7

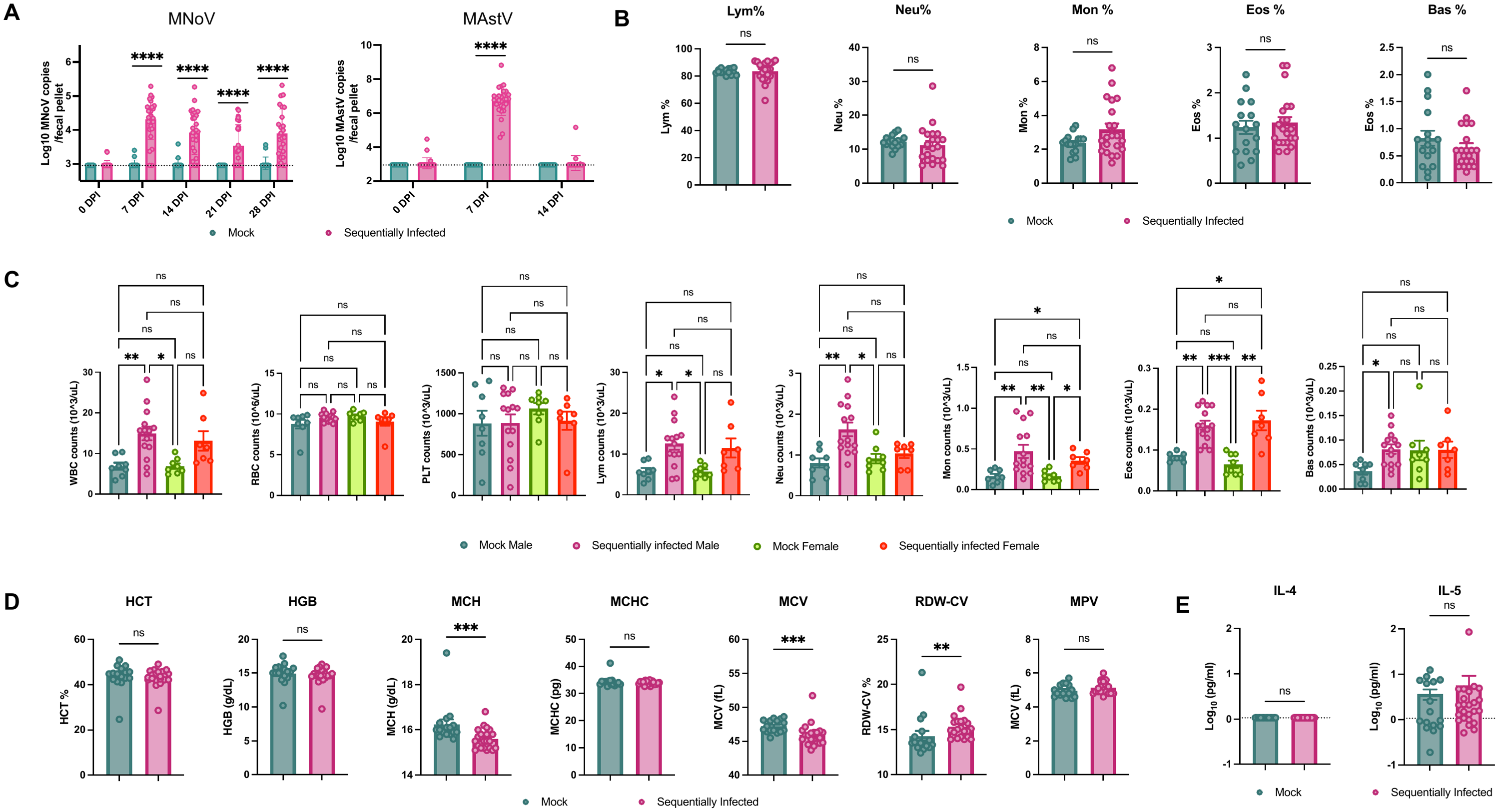
·

**Supplemental Figure 1. Sequential infection does not alter relative proportions of white blood cells.**

(**A**) Murine norovirus strain CR6 (MNV) and murine astrovirus (MAstV) genome copies detected in fecal pellets of mock- (n = 18) and sequentially-infected (n = 25) mice at indicated timepoints post-inoculation.

(**B**) Frequency of lymphocytes (Lym), neutrophils (Neu), monocytes (Mon), eosinophils (Eos), and basophils (Bas) in the hematological analysis of mock- (n = 16) and sequentially-infected (n = 21) mice at 10 weeks of age.

(**C**) Separation of CBC data by sex for mock- and sequentially-infected mice.

(**D**) Hematological analysis of hematocrit (Hct), hemoglobin (Hgb), mean corpuscular hemoglobin (MCH), mean corpuscular hemoglobin concentration (MCHC), mean corpuscular volume (MCV), mean platelet volume (MPV) and increased red cell distribution width (RDW-CV).

(**E**) Serum cytokines IL-4 and IL-5 in mock (n = 16) and sequentially-infected (n = 21) mice at 10 weeks of age.

Columns show median values, error bars represent the standard error of the mean, and dotted lines indicate the LOD of the assays. Data are representative of 3 technical replicates. The undetectable samples were given a value of LOD. In A, significance was determined using a two-way ANOVA test with the Geisser_Greenhouse correction; In B, D, and E, significance was determined using unpaired Mann-Whitney test; In C, significance was determined using Dunn’s multiple comparisons correct test: *p < 0.05;**p < 0.01; ***p < 0.001; ****p < 0.0001; ns, not significant.

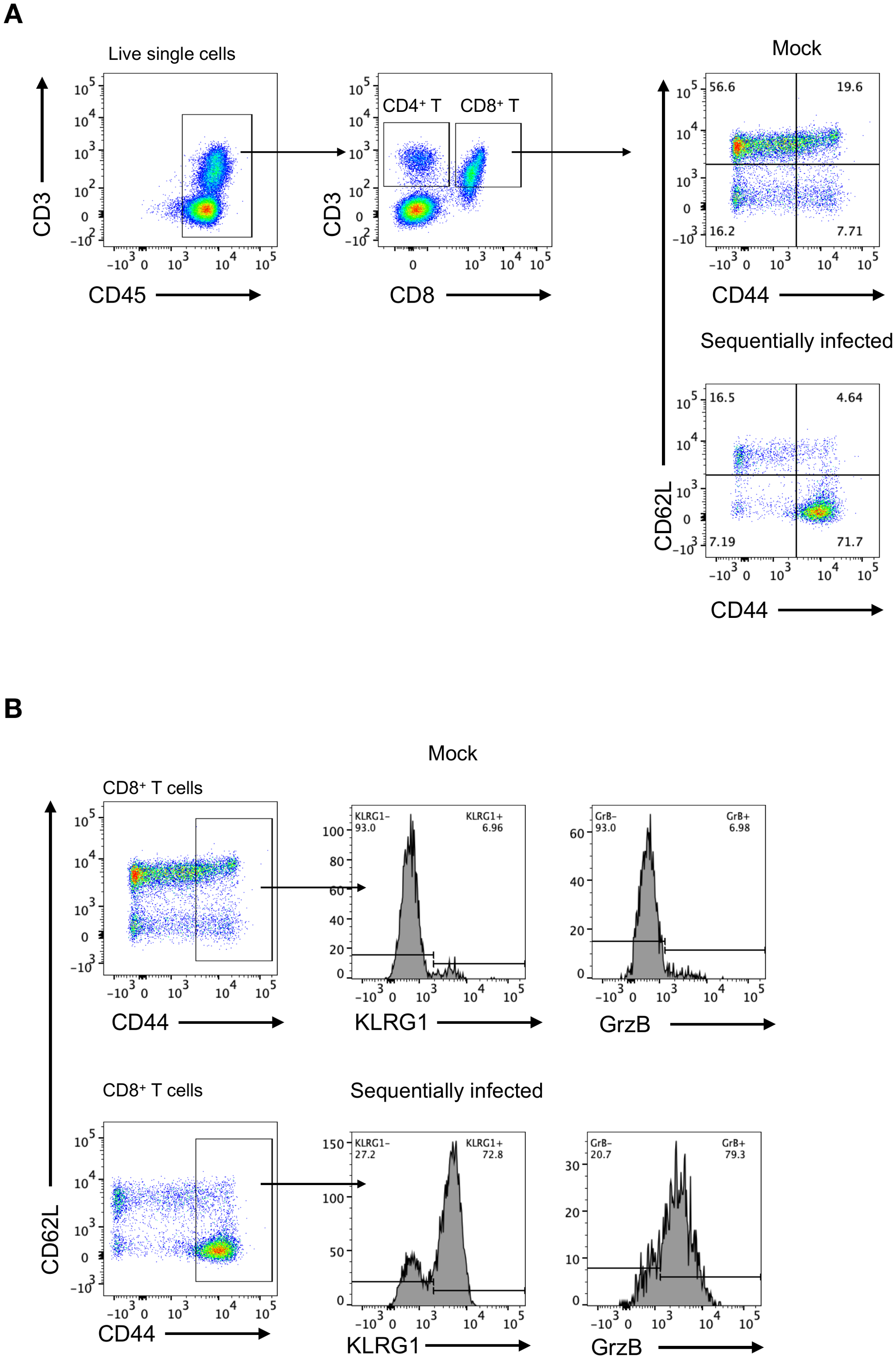

**Supplemental Figure 2. Gating strategy for analysis of T cells.**

(**A**) T cells from mock- (n = 16) or sequentially-infected (n = 21) mice were analyzed at 10 weeks of age. PBMC live single cells were gated for CD45+ followed by CD3+CD4+ or CD3+CD8+ cell populations. CD44^lo^/CD62L^hi^ (Naive), CD44^hi^/CD62L^hi^ (antigen-experienced central memory, CM), and CD44^hi^/CD62L^lo^ (antigen-experienced effector memory, EM) CD8^+^ T cells were evaluated with CD44 and CD62L expression.

(**B**) Gating of KLRG1+ and GrzB+ cells from CD44^hi^ (antigen-experienced) CD8^+^ T cells from PBMCs of mock- or sequentially-infected mice.

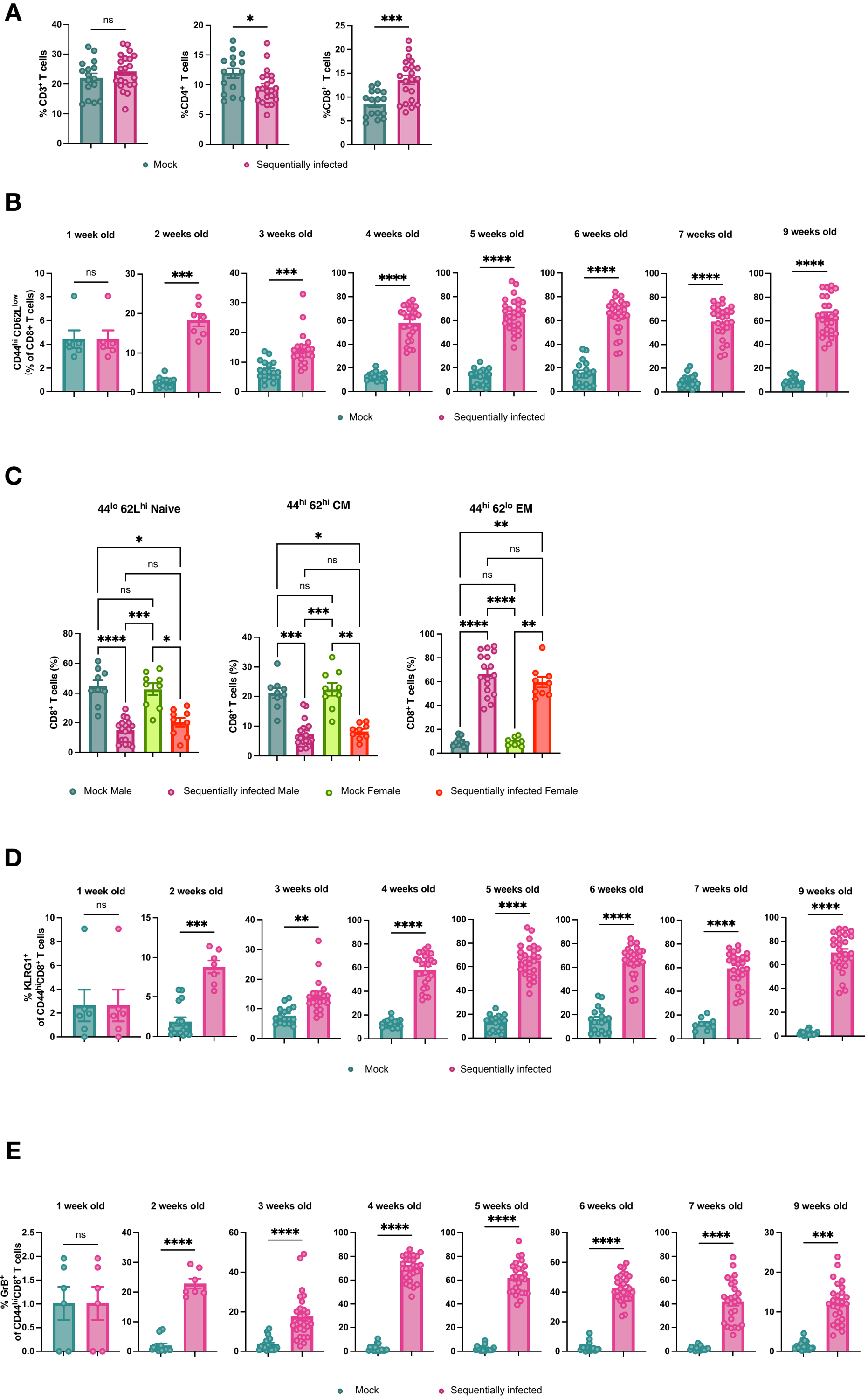

**Supplemental Figure 3. PBMC T cell subsets vary between mock- and** **sequentially-infected mice.**

(**A**) Frequency of PBMC CD3^+^, CD4^+^, and CD8^+^ T cells from mock- (n = 16) or sequentially-infected (n = 21) mice at 10 weeks of age.

(**B**) Frequency of CD44^hi^ (antigen-experienced) CD8^+^ T cells from PBMCs of mock- or sequentially-infected mice between 1 week and 9 weeks of age.

(**C**) CD44^lo^/CD62L^hi^ (Naive), CD44^hi^/CD62L^hi^ (antigen-experienced central memory, CM), and CD44^hi^/CD62L^lo^ (antigen-experienced effector memory, EM) CD8^+^ T cells of mock-infected and sequentially infected mice, separated by sex.

(**D**) Frequency of KLRG^+^ CD44^hi^ CD8^+^ T cells from PBMCs of mock- or sequentially-infected mice between 1 week and 9 weeks of age.

(**E**) Frequency of GrzB^+^ CD44^hi^ CD8^+^ T cells from PBMCs of mock- or sequentially-infected mice between 1 week and 9 weeks of age.

Columns show median values, error bars represent the standard error of the mean. In A, significance was determined using unpaired Mann-Whitney test; In B, D and E, significance was determined using a two-way ANOVA test with the Geisser_Greenhouse correction; In C, significance was determined using Dunn’s multiple comparisons correct test: *p < 0.05; **p < 0.01; ***p < 0.001; ****p < 0.0001; ns, not significant.

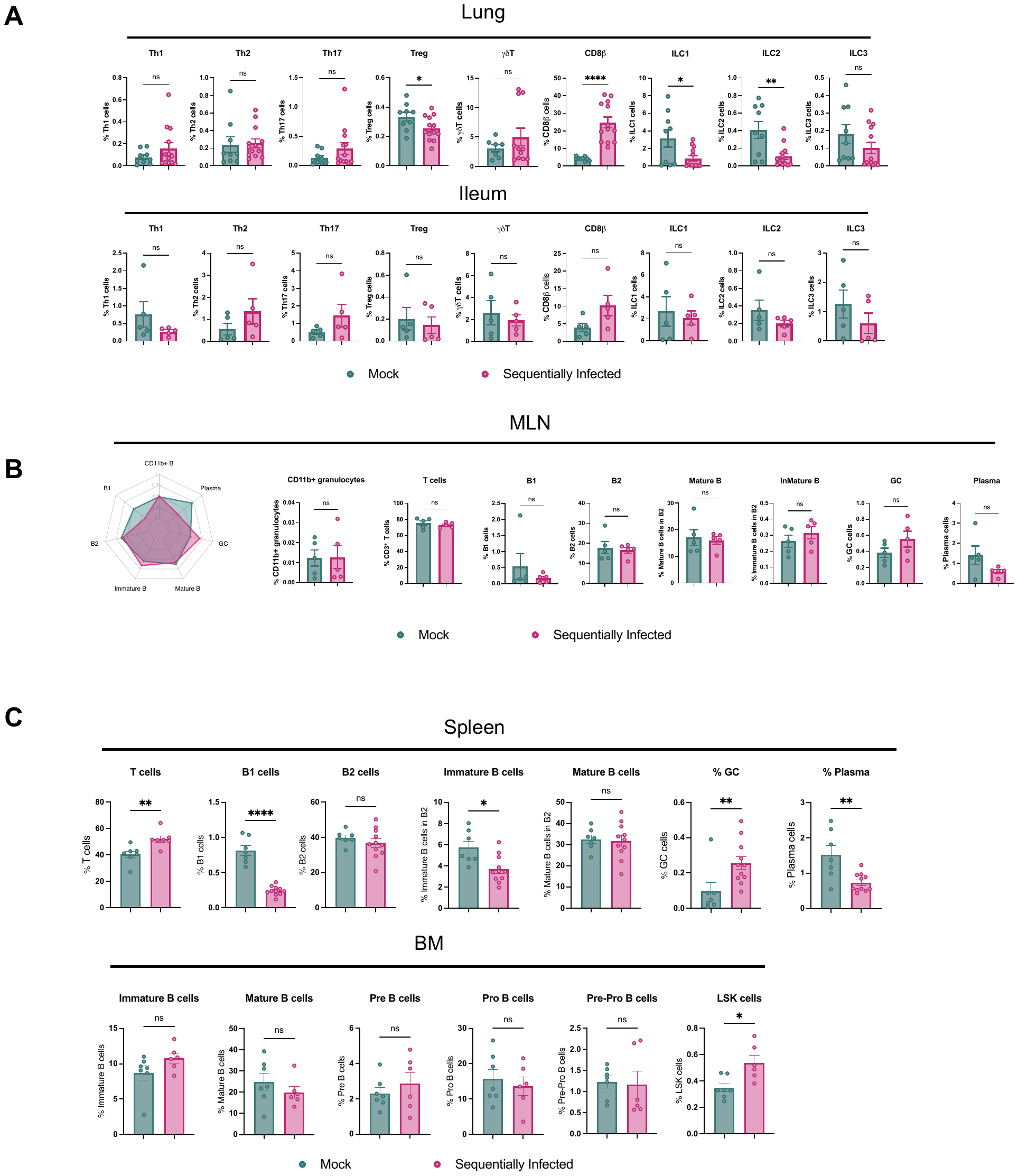

**Supplemental Figure 4. Sequential viral exposure shapes immune cell profiles in tissues.**

(**A**) Frequency of ILC, innate lymphoid cells; T_h_, CD4 T helper cells; T_reg_, regulatory CD4 T cells; γδ T, gamma delta T cell in the lung and ileum of mock- and sequentially-infected mice at 10 weeks of age.

(**B**) Immune cell types isolated and enumerated by flow cytometry from the mesenteric lymph node (MLN) of mock- and sequentially-infected mice, including granulocytes, T cells, B1 cells, B2 cells, mature B cells, immature B cells, GC and plasma cells.

(**C**) Frequency of T cells, B1 cells, B2 cells, mature B cells, immature B cells, GC and plasma cells from the spleen as well as the mature B cells, immature B cells, Pre B cells, Pro B cells, Pre-pro B cells, and LSK cells from bone marrow (BM) of mock- and sequentially-infected mice at 10 weeks of age. Columns show median values, error bars represent the standard error of the mean. Significance was determined using unpaired Mann-Whitney test: *p < 0.05; **p < 0.01; ***p < 0.001; ****p < 0.0001; ns, not significant.

**Supplemental Figure 5. Gating strategy for tissue immune cell profiling.**

Ten-week-old mock- and sequentially-infected mice were used to analyze immune cell profiles in different tissues. (**A**) Lung cells were gated for lymphocytes (FSC-A/SSC-A), singlets (SSC-W/SSC-H), live cells (Aqua-), CD45+ followed by CD3+ or NK1.1+ cell populations to identify ILC, innate lymphoid cells; T_h_, CD4 T helper cells; T_reg_, regulatory CD4 T cells; γδ T, gamma delta T cells. Single positive staining for T-bet, Gata3, and Rorγt were used to define Th1, Th2, and Th17 lineages, respectively.

(**B**) Gating of CD3+ T cells, CD11b+ granulocytes, B1, mature or immature B cells, germinal center (GC) B cells and plasma cells from splenocytes of mock- or sequentially-infected mice.

(**C**) Gating of Lin^-^Sca1^+^c-Kit^+^ (LSK), mature or immature B cells, Pre B cells, Pro B cells, and Pre- Pro B cells from bone marrow of mock- or sequentially-infected mice.

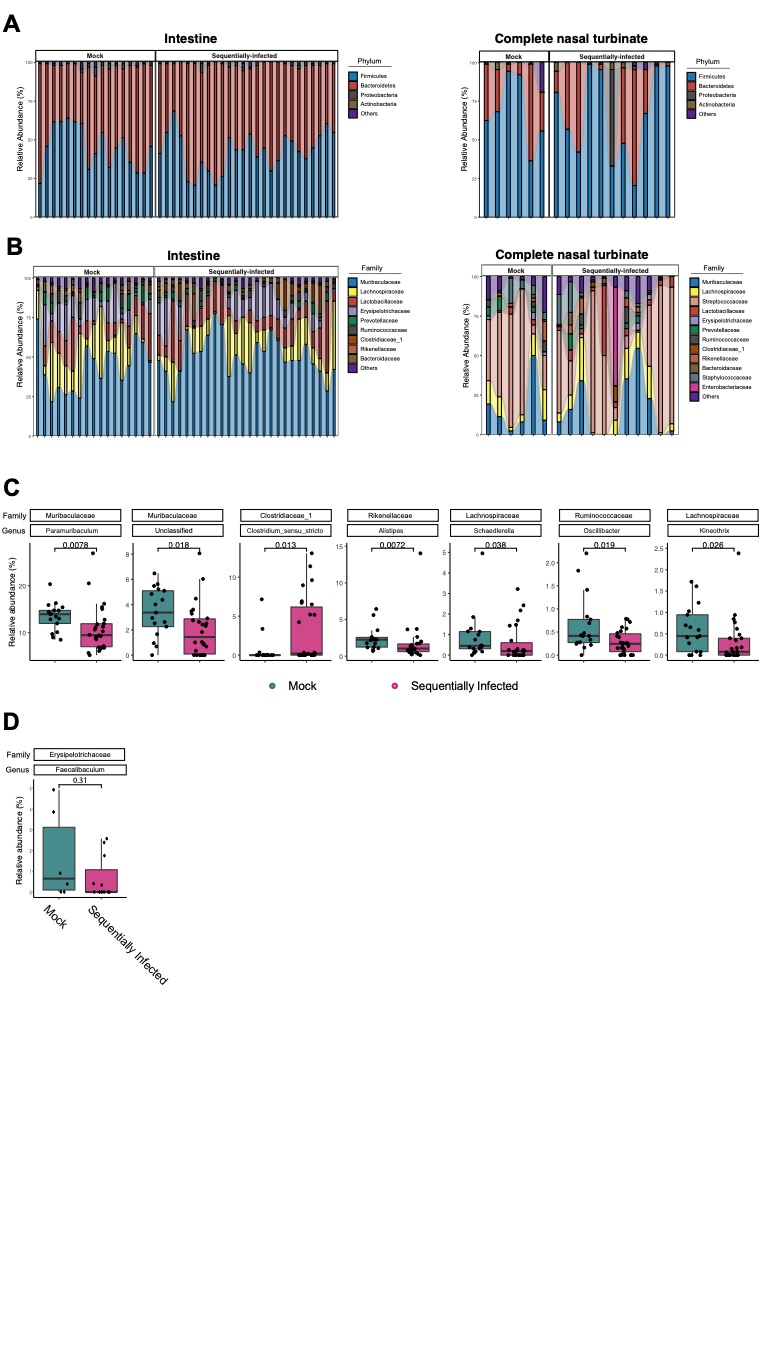

**Supplemental Figure 6.** **Comparison of intestinal and complete nasal turbinate microbiota between mock- and sequentially-infected mice.**

(**A, B**) Intestine (mock-: n = 17, sequentially-infected: n = 26) and complete nasal turbinate (mock-: n = 6, sequentially-infected: n = 11) microbiota composition in mock- and sequentially-infected mice at (**A**) phylum and (**B**) species levels.

(**C, D**) Relative abundance of potential discriminant taxa from linear discriminant analysis within the (**C**) intestinal and (**D**) complete nasal turbinate at the genus level.

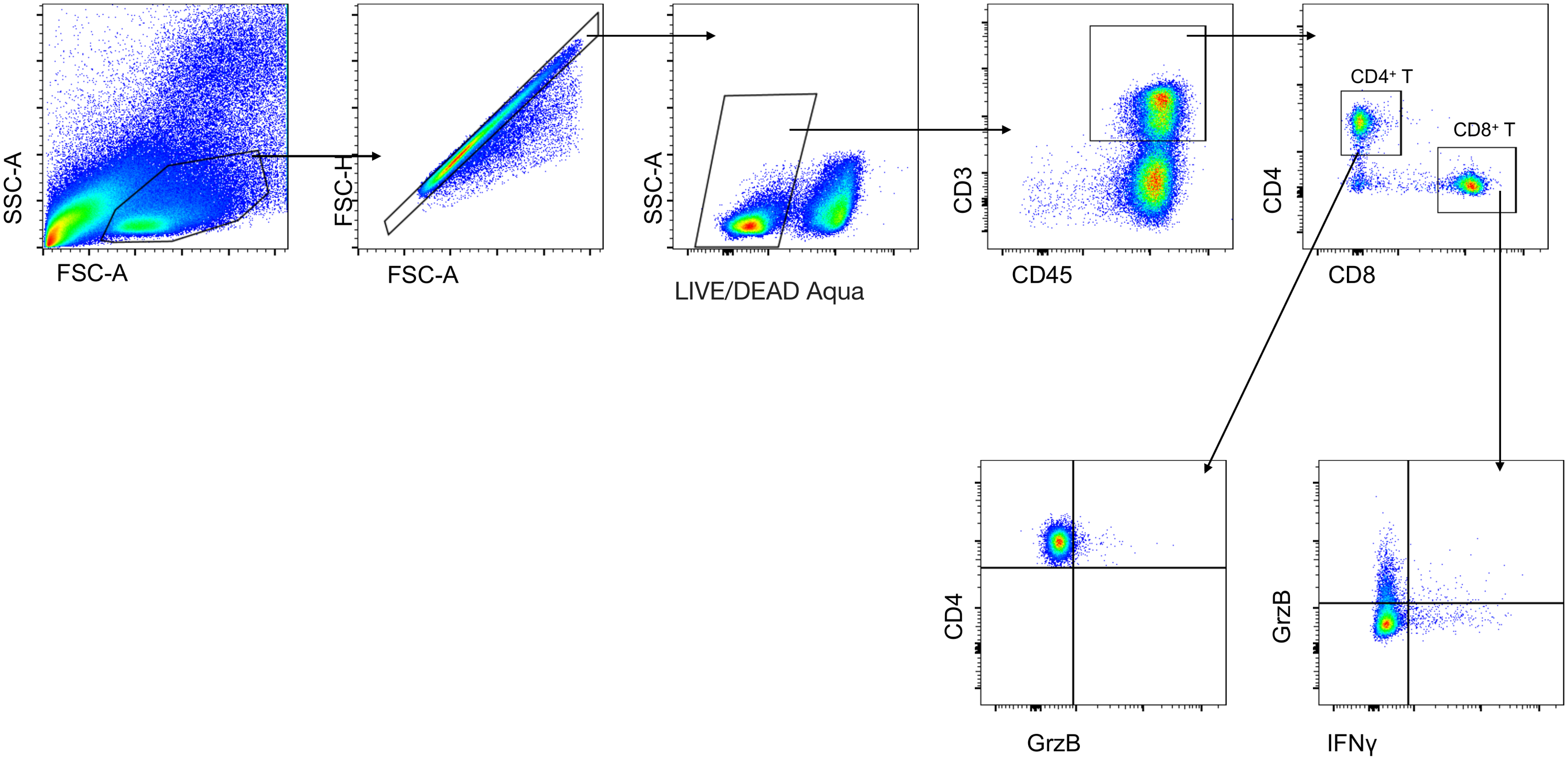

**Supplemental Figure 7. Gating strategy for post-immunization T cell responses.**

Ten-week-old mock- and sequentially-infected mice were immunized with ChAd-SARS-CoV-2-S. T cell responses were analyzed in splenocytes at 5 weeks post-vaccination. Cells were gated for lymphocytes (FSC-A/SSC-A), singlets (SSC-W/SSC-H), live cells (Aqua-), CD45+, CD19- followed by CD4+ or CD8+ cell populations expressing IFNγ or granzyme B.
